# ATAC-seq identifies chromatin landscapes linked to the regulation of oxidative stress in the human fungal pathogen *Candida albicans*

**DOI:** 10.1101/2020.05.07.080739

**Authors:** Sabrina Jenull, Michael Tscherner, Theresia Mair, Karl Kuchler

**Author notes:** contributed equally (Sabrina Jenull, Michael Tscherner).

## Abstract

Human fungal pathogens often encounter fungicidal stress conditions upon host invasion, but they can swiftly adapt by transcriptional reprogramming that enables pathogen survival. Fungal immune evasion is tightly connected to chromatin regulation. Hence, fungal chromatin modifiers pose alternative treatment options to combat fungal infections. Here, we present an ATAC-seq protocol adapted for the opportunistic pathogen *Candida albicans* to gain further insight into the interplay of chromatin accessibility and gene expression mounted during fungal adaptation to oxidative stress. The ATAC-seq workflow facilitates the robust detection of genomic regions with accessible chromatin, but also allows for the precise modeling of nucleosome positions in *C. albcians*. Importantly, the data reveal genes with altered chromatin accessibility in upstream regulatory regions, which correlate with transcriptional regulation during the oxidative stress response. Interestingly, many genes show increased chromatin accessibility yet no change in gene expression upon stress exposure. Such chromatin signatures could predict yet unknown regulatory factors under highly dynamic transcriptional control. In addition, *de novo* motif analysis in genomic regions with increased chromatin accessibility upon hydrogen peroxide treatment shows significant enrichment for Cap1 binding sites, a major factor of oxidative stress responses in *C. albicans*. Taken together, the ATAC-seq workflow enables the identification of chromatin signatures and uncovers the dynamics of regulatory mechanisms mediating environmental adaptation of *C. albicans* to host immune surveillance.

**Importance:** The opportunistic fungal pathogen *Candida albicans* colonizes and infects various tissues and organs of the human host. This is due to its rapid environmental adaptation facilitated by changes in gene expression coupled to chromatin alterations. Recent advances in chromatin profiling approaches, such as the development of ATAC-seq, shed light on the dynamic interplay of chromatin accessibility and transcriptional control. The here presented expansion of the ATAC-seq method to *C. albicans* demonstrates the robustness of ATAC-seq to detect dynamic modulations of chromatin accessibility in response to oxidative stress. This work serves as a basis to further exploit this application to characterize regulatory mechanisms that drive fungal environmental adaptation, such as during host invasion, and thus, will open novel antifungal treatment strategies targeting fungal chromatin regulation.

## Introduction

Human fungal pathogens respond to host immune defense through numerous mechanisms, including chromatin-mediated adaptive gene expression ^1–3^. For example, immune defense or environmental changes can trigger pathogen responses through extracellular sensing, intracellular signal transduction and transcriptional reprogramming ^4,5^. Transcriptional changes require a tight interplay of chromatin states and transcription factors ^6,7^, as the swift adaptation to environmental changes is often paramount for a successful lifestyle or immune evasion. For instance, pathogens encounter a number of extreme stress conditions during the course of an infection. These include limitations in nutrient availability and the cytotoxic attack by the host immune system ^8,9^. The opportunistic fungal pathogen *Candida albicans* is an extraordinary example of how a pathogen can occupy multiple host niches to persist and survive. *C. albicans* lives as a harmless commensal in the majority of humans, colonizing mucosal surfaces and epithelial barriers, especially the intestinal tract. However, severe immunodeficiency or a dysregulation of the host microbiota, turns *C. albicans* into an invasive pathogen that can infect virtually any tissue or organ in the human body ^10,11^. In the last decade, numerous efforts have been made to better understand fungal pathogenicity mechanisms, including the transcriptional regulation of environmental adaptation and virulence factors, such as the switch between different cellular morphologies ^12–14^. Given the pivotal interplay of chromatin modifications and transcription control, it is hence not surprising that several *C. albicans* chromatin-modifying factors play important roles in fungal virulence. For instance, the function of the histone acetyl transferases (HATs) Gcn5, Hat1 and Rtt109 are crucial for morphogenesis and virulence ^15–19^. Likewise, the histone deacetylase (HDAC) complex Set3C controls transcriptional kinetics during the morphological transition from yeast growth to filamentous hyphal growth. Remarkably, genetic ablation of *SET3* abrogates fungal virulence ^20^. Hence, attacking chromatin modifiers provides a new option for antifungal drug development. This requires immediate attention in clinical settings, given the rapid emergence of antifungal drug resistance in Candida spp such as *C. glabrata* or *C. auris* ^17,21–25^. However, a better understanding of the interplay between chromatin architecture in the pathogen and transcriptional reprogramming during host interaction would further aid discovery of new antifungals targeting chromatin function.

Several methods including chromatin immunoprecipitation (ChIP), micrococcal nuclease (MNase) digestion of chromatin coupled to next generation sequencing (ChIP-seq and MNase-seq, respectively) or DNase-seq have been employed to analyze chromatin accessibility and nucleosome positioning to reveal regulatory mechanisms ^26–30^. Recently, the assay for transposable-accessible chromatin using sequencing (ATAC-seq) emerged to probe for native chromatin states ^30^, which includes accessibility as well as nucleosome positioning ^31^. Due to its technical simplicity and the low sample input requirements, ATAC-seq has been widely applied for chromatin profiling in various cell types ^31–34^, tissues ^35,36^ and even single cells ^37,38^. Moreover, it has proved as a useful tool for identifying sequence motifs decorated by transcriptional regulators and for predicting gene transcription ^39–42^.

Here, we aimed to adapt the original ATAC-seq protocol ^30^ for the human fungal pathogen *C. albicans*. We developed a modified protocol and bioinformatics workflow that enables the sensitive detection of changes in the global chromatin landscapes in response to environmental stress. We have chosen oxidative stress as environmental cue, because it triggers genome-wide changes in gene expression ^14,16^ and because it closely mimics the oxidative immune defense fungal pathogens face during host invasion ^43,44^. Moreover, transcriptomics data for *C. albicans* challenged with hydrogen peroxide (H_2_O_2_) and genome-wide binding data of the oxidative stress transcriptional regulator Cap1 are available ^16,45^. Hence, we combine these data sets with the present ATAC-seq data for *C. albicans*. With this approach, we demonstrate that H_2_O_2_-treatment of fungal cells increases chromatin accessibility in upstream regions of genes associated with the oxidative response, and we show that those genes tend to be transcriptionally upregulated. Moreover, signatures of accessible chromatin regions enable the prediction of putative novel regulators of stress signaling that are not yet linked to transcriptional control. In addition, genomic regions with elevated ATAC-seq signal are enriched in binding sites for the key regulator Cap1, demonstrating the potential for *de novo* motif discovery of regulatory factors. In summary, we show the versatility of ATAC-seq chromatin profiling in *C. albicans*, especially when combined with complementary next generation sequencing data such as RNA-seq. This approach uncovers dynamic and complex regulatory mechanisms during environmental adaptation of pathogens.

## Results & Discussion

### ATAC-seq in *C. albicans* reflects nucleosomal organization genome-wide

We adopted and optimized an ATAC-seq method originally developed for the fungi *S. cerevisiae* and *Schizosaccharomyces pombe* ^31^, so as to study alterations in chromatin landscapes upon oxidative stress in the human fungal pathogen *C. albicans*. Therefore, logarithmically growing *C. albicans* cells were either treated with 1.6 mM H_2_O_2_ for 15 minutes in YPD or left untreated. After spheroblasting, cells were then subjected to sample preparation for ATAC-seq analysis. To correct for the known sequence bias of the Tn5 transposase ^46^, naked genomic DNA (gDNA) prepared from YPD-grown *C. albicans* was included into the workflow (Figure 1A; see Materials & Methods for details). Three biological replicates were prepared for each condition. Analysis of PCR-amplified ATAC-seq libraries from intact fungal cell nuclei (chromatin) showed a distinct fragment size distribution reminiscent of nucleosomal periodicity. Importantly, ATAC-seq libraries prepared from naked gDNA had a less complex fragment distribution as they lacked nucleosomal patterns (Figure 1B). The same trend was observed after 75 bp paired-end sequencing (PES), where the fragment length distribution of PES reads from fungal chromatin showed a predominant peak of shorter fragments around < 150 bp, representing putative nucleosome-free regions. Another peak with an average fragment length of 200 bp reflected DNA regions occupied by one nucleosome (Figure 1C; ^47^). Additionally, a weak enrichment for di-nucleosomal peaks was detected. Again, ATAC-seq libraries from gDNA did not reveal nucleosomal presence (Figure 1C). Instead, the majority of PES read fragments were below 150 bp, which was consistent with fragment length distribution of Tn5-generated DNA sequencing libraries ^48^. Notably, the number of mapped reads were evenly distributed among the *C. albicans* chromosomes, thus reflecting total chromosome sizes and suggesting no chromosomal bias (Figure S1A-B). Inspection of ATAC-seq reads aligned to the *C. albicans* genome further revealed a distinct read coverage profile of accessible chromatin regions in nuclear chromatin purified from H_2_O_2_-treated (H_2_O_2_) and non-treated (YPD) cells, which was not apparent in gDNA libraries (Figure 1D shows a region of chromosome 1 as an example).

**Figure 1.**
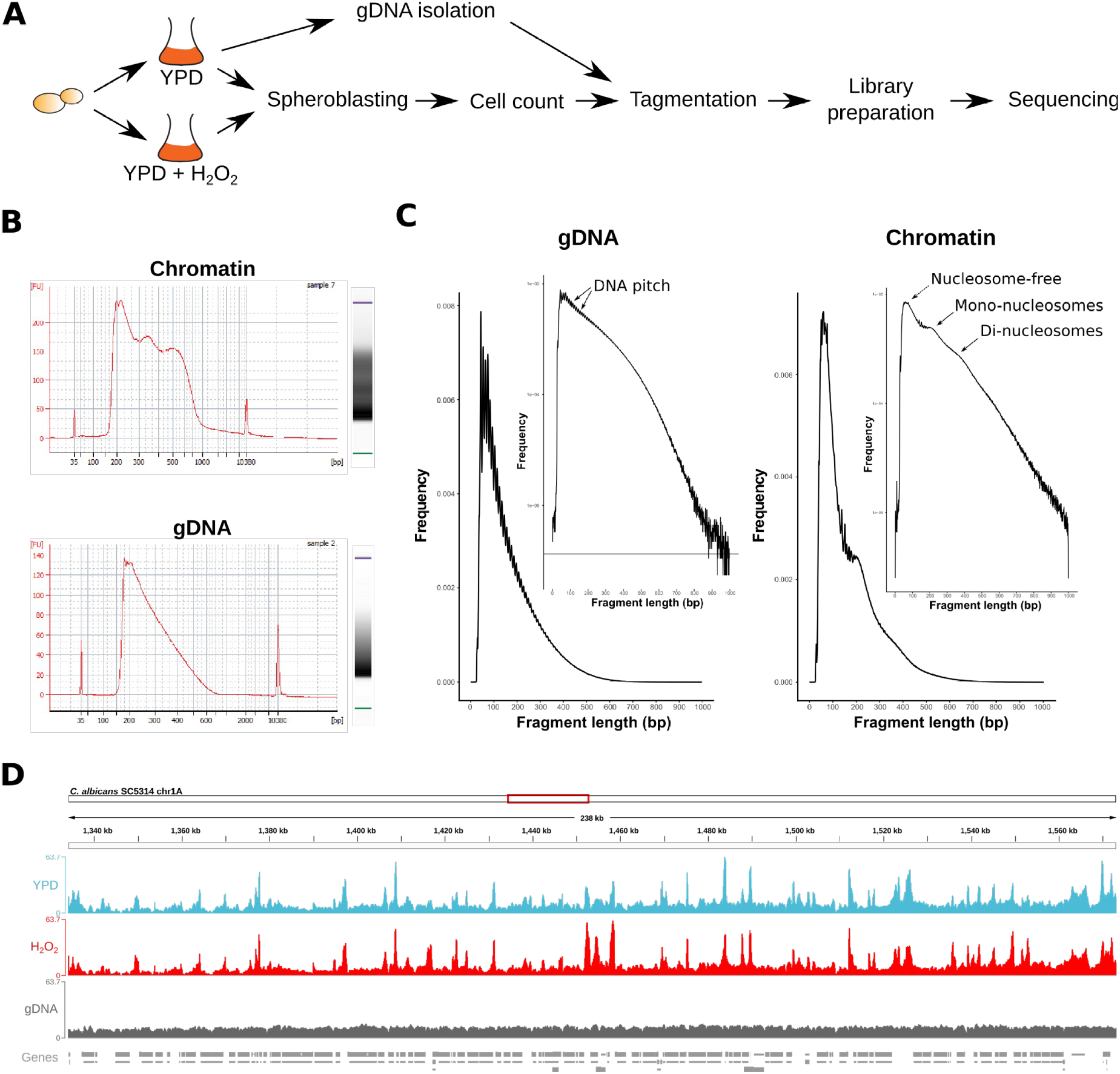
Quality control of ATAC-seq libraries from *C. albicans* treated or non-treated with H_2_O_2_. **A**) Experimental set-up for ATAC-seq profiling of 5×10^6^ *C. albicans* (see Materials & Methods for details). **B**) Bioanalyzer electrophoretic profiles and gel image from ATAC-seq libraries prepared from fungal chromatin (isolated nuclei) or naked gDNA. The x-axis from the electropherogram depicts the fragment size distribution (bp) of tagmented samples and the y-axis represents the abundance. A distinct nucleosomal pattern (mono-, di- and tri-nucleosomes) is visible in chromatin samples, but not in tagmented gDNA samples. **C**) Fragment length distribution of ATAC-seq reads from gDNA, nuclei prepared from H_2_O_2_-treated cells (H_2_O_2_) and non-treated cells (YPD). The fragment length in bp (x-axis) from one representative biological replicate from each group is plotted against its frequency (y-axis). The graph insert shows the log-transformed histogram. Signals originating from the DNA helical pitch ^31^ are emphasized by arrows on the gDNA plot. Arrows on the chromatin plot indicate nucleosome-free ATAC-seq read fragments and nucleosome occupied read fragments. **D**) IGV browser snap-shot from a 238 kb region of the *C. albicans* chromosome 1A of ATAC-seq reads from aligned bam files. The biological replicates from each sample group were pooled. Genes are depicted as grey boxes below the sequencing read coverage tracks.

To further test the robustness of our ATAC-seq workflow, we first selected ATAC-seq PES read fragments below 100bp, which corresponded to putative nucleosome-free genomic regions ^30^. These, data were subjected to nucleosomal occupancy prediction using the NucleoATAC tool ^31^. First, read coverage profiles of nucleosome-free ATAC-seq peaks and NucleoATAC-called nucleosomes were inspected at an example locus on chromosome 6 containing four open reading frames (ORFs) (Figure 2A). Pronounced nucleosome-free ATAC-seq peaks, flanked by nucleosomal signals as predicted by NucleoATAC, were detected in upstream promoter regions (Figure 2A, black arrows). Moreover, nucleosomal positions predicted by NucleoATAC correlated very well with published *C. albicans* MNase-seq data ^49^ (Figure 2B). By averaging the read coverage of nucleosome-free and mononucleosmal ATAC-seq signals across all *C. albicans* transcripts relative to the transcription start site (TSS), we further observed an enrichment of nucleosome-free ATAC-seq reads adjacent to the TSS. Mononucleosomal ATAC-seq signals were enriched up- and downstream of the nucleosome-free peak, reflecting a well-positioned +1 nucleosome and a typical nucleosomal organization of canonical active gene promoters ^26,50^ (Figure 2B). Taken together, ATAC-seq libraries generated from *C. albicans* native chromatin share typical features of mammalian and other fungal (*S. cerevisiae* and *S. pombe*) ATAC-seq samples ^30,31^. Hence, the approach is suitable to probe chromatin accessibility and nucleosomal positioning in *C. albicans* and their changes during stress response.

**Figure 2.**
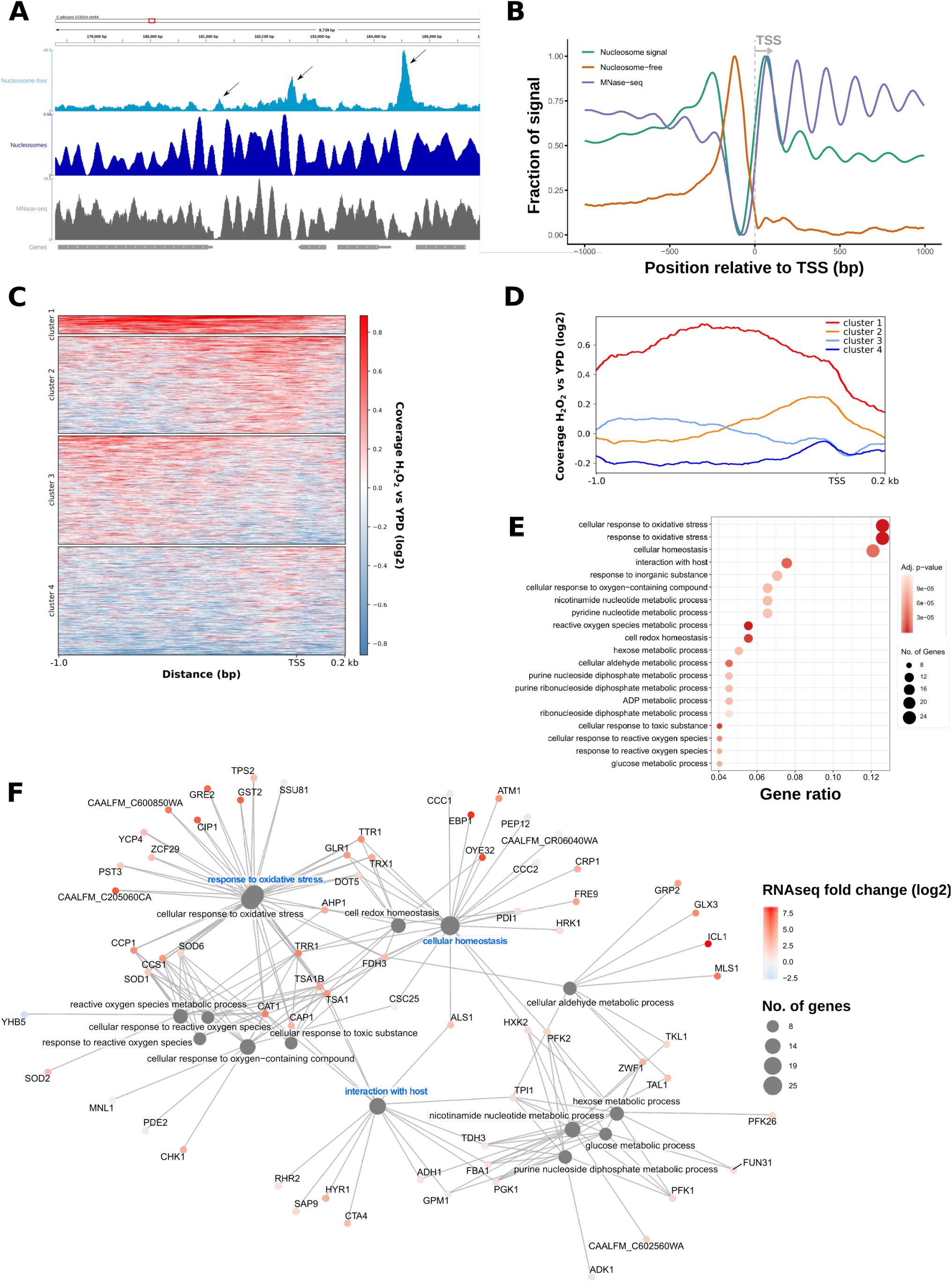
Chromatin accessibility is increased at promoters associated with oxidative stress genes. **A-B**) ATAC-seq analysis in *C. albicans* allows for nucleosomal occupancy prediction. Comparison of coverage profiles of nucleosome-free ATAC-seq read signals with mono-nucleosomal position prediction and MNase-seq data ^49^ at a region on chromosome 6 (A) or across all *C. albicans* transcriptional start sites (+/- 1000 bp relative to TSS) (B). Data represent ATAC-seq samples from YPD grown cells and pooled biological replicates (see Materials & Methods for details). A distinct +1 nucleosome signal is readily detected which is preceded by a nucleosome-free region upstream of the TSS. In panel A, genes are depicted as grey boxes below the sequencing read coverage tracks and white arrows within the box indicate the direction of transcription. The black arrows in the top IGV track indicate nucleosome-free ATAC-seq signals flanked by nucleosomes. **C-D**) Differential ATAC-seq nucleosome-free read coverage (log2 fold change) of H_2_O_2_-treated (H_2_O_2_) vs nontreated (YPD) cells clustered with k-means clustering and represented as heatmaps (C) or read coverage profiles across all *C. albicans* promoters (−1000bp/+200bp relative to the TSS). **E-F**) GO term analysis of cluster 1 from panel C-D represented as dotplot (E) and cnet plot (F). Genomic regions represented in cluster 1 were annotated to the next downstream gene, which were subjected to GO term analysis. The y-axis of the dotplot shows the name of the GO term and the x-axis indicates the genes enriched in this GO term relative to all detected genes with increased chromatin accessibility upon H_2_O_2_ stress. The color scale indicates the adjusted p-value of the GO enrichment analysis. The dot size represents the number of genes from cluster 1 in each GO term. The cnet plot shows each GO term category and the associated genes enriched in this GO term. Gene dots are overlaid with a colour gradient representing the log2 fold change in transcription (from RNA-seq data) in response to oxidative stress of the depicted genes.

### ATAC-seq detects genome-wide changes in chromatin accessibility after oxidative stress

Next, we aimed to assess whether ATAC-seq in *C. albicans* is able to measure changes in chromatin accessibility at genomic regions associated with the oxidative stress response. Therefore, we analyzed the differential read coverage of nucleosome-free ATAC-seq peaks in H_2_O_2_-treated and nontreated cells. Then, we further clustered all *C. albicans* promoter regions (−1000/+200bp upstream of the TSS) based on their differential read coverage profile using *k-means* clustering (Figure 2C). Genomic regions with a moderate increase in nucleosome-free ATAC-seq peak signals around the TSS were enriched in cluster 2, while cluster 3 showed a subtle increase in ATAC-seq read coverage more distal relative to the TSS (Figure 2C-D, orange and light blue lines, respectively, in panel D). The most striking increase in chromatin accessibility (i.e. increased nucleosome-free ATAC-seq reads) proximal to the TSS, as well as at least 1 kb upstream of the TSS, was observed in response to oxidative stress for loci contained in cluster 1 (Figure 2C-D, red line in panel D)(Figure 2C-D). GO term enrichment analysis of promoter-associated genes from cluster 1 showed that around 12 % of genes in cluster 1 are related to the oxidative stress response (Figure 2E). Cluster 4 included regions with decreased chromatin accessibility upstream of the TSS in response to H_2_O_2_ treatment (Figure 2C-D, dark blue lines in panel D). Genes from this cluster were involved in ribosome- and translation-related processes (Figure S2A), most of which are often downregulated during fungal stress adaptation ^51–55^.

The degree of chromatin accessibility is typical for genomic regions experiencing distinct transcriptional activities. For instance, the TSS of active gene promoters and enhancers are associated with increased chromatin accessibility relative to inactive elements and heterochromatic regions ^28,56–58^. Hence, the combination of ATAC-seq and RNA-seq can provide highly useful insights into the chronological dynamics of changing chromatin accessibility and subsequent transcriptional responses ^42,59^. In a previous study, we performed RNA-seq analysis in *C. albicans* in response to H_2_O_2_ ^16^. Since the conditions were almost identical (see Material & Methods for details), we used this data set to test how the enrichment of nucleosome-free ATAC-seq read signals correlated with transcript abundance in *C. albicans*. When assessing the log2-fold change from our RNA-seq data set of genes with increased chromatin accessibility upon H_2_O_2_ treatment in upstream genomic regions (cluster 1), we observed that transcription of the majority of cluster 1 genes was also upregulated in response to H_2_O_2_ (Figure 2F). For instance, *CAT1*, which encodes the H_2_O_2_-detoxifying catalase ^60^ as well as the targets of the oxidative stress-related Cap1 transcription factor *OYE32, CIP1* and *EBP1^61^*, were among genes showing remarkably increased chromatin accessibility and transcriptional induction in H_2_O_2_-treated cells (Figure 2F). In addition, *ICL1* and *MSL1* encoding for glyoxylate cycle enzymes ^62^ showed increased ATAC-seq peak signals, reflecting their robust transcriptional induction upon oxidative stress (Figure 2F). Similarly, genes with decreased ATAC-seq read coverage upstream of their TSS were likewise repressed upon stress (Figure S2B). Hence, changes in ATAC-seq read coverage corresponded to alterations in gene expression.

To further dissect significant changes in chromatin accessibility, nucleosome-free ATAC-seq signals were subjected to peak calling with ATAC-seq libraries from gDNA as background noise control and quantification of called peaks (see Material & Methods for details). The genomic locations of the majority of called nucleosome-free ATAC-seq peaks were located within gene promoters, exons and 5’ untranslated region (UTR) regions (Figure 3A) reflecting a usual distribution of genomic features among called ATAC-seq peaks ^63^. Of note, substantial peak overlaps between these three genomic regions were observed, indicating peaks stretching from promoter regions into the 5’ UTRs and the first exon. Interestingly, a previous study observed ATAC-seq signals accumulating towards sub-telomeric regions ^39^. Thus, we further inspected the genome-wide distribution of differentially enriched peaks, but we did not detect a bias towards specific chromosomal regions including telomeres (Figure 3B).

**Figure 3.**
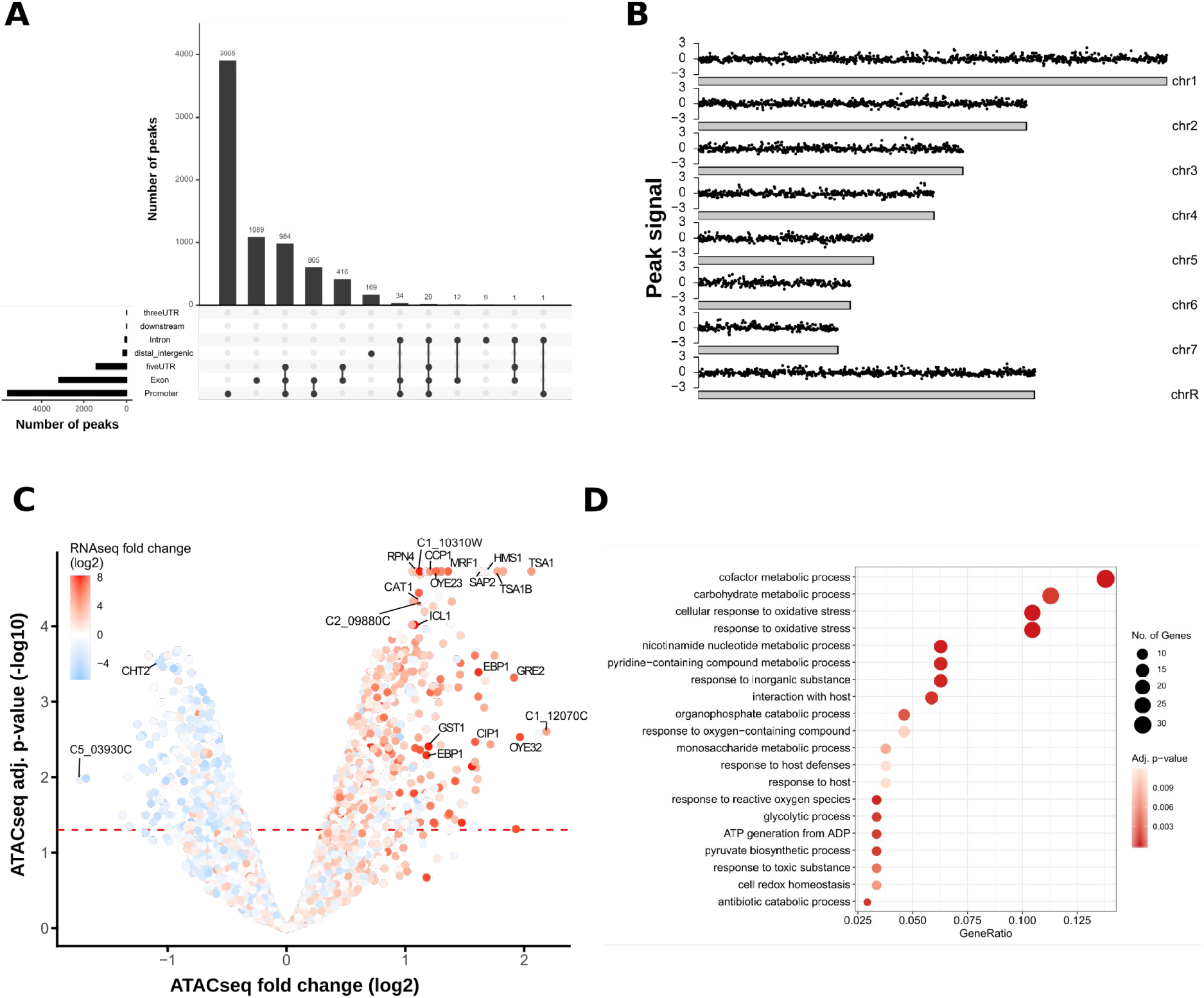
ATAC-seq nucleosome-free peak annotation and differential ATAC-seq peak analysis. **A**) The majority of called ATAC-seq nucleosome-free peaks are within gene promoters. The vertical bars of the UpSet plot represent the number of peaks spanning one or more specific genomic regions (overlaps between genomic regions are indicated by connection lines between the dots). For instance, the first bar indicates the number of peaks occurring exclusively in promoters, while the third bar represents peaks spanning promoter, 5’UTR and exonic regions. The horizontal bars show the total number of peaks for each genomic annotation, independent of overlaps. **B**) Karyoplots showing the genomic peak location on each chromosome of differential ATAC-seq nucleosome-free read signals (log2 fold change) of H_2_O_2_-treated vs non-treated cells. The log2 fold change is depicted on the y-axis and the x-axis represents each chromosome of the *C. albicans* genome. **C**) Increased ATAC-seq nucleosome-free peak signals correlate with increased transcript levels. Volcano plot depicting the differential peak signal analysis of nucleosome-free ATAC-seq peak signals between H_2_O_2_-treated and non-treated cells. The y-axis represents the negative log10 adjusted p-value (FDR) and the x-axis depicts the log2 fold change H_2_O_2_ treatment vs no treatment. The horizontal dashed red line indicates a FDR of 0.05. Each dot represents one peak annotated to the next downstream gene which is overlaid with a colour gradient indicating the log2 fold change (H_2_O_2_ treatment vs no treatment) from RNA-seq data. **D**) Peaks with increased ATAC-seq nucleosome-free signals in H_2_O_2_-treated cells are upstream of genes related to oxidative stress response. The y-axis shows the GO term name and the x-axis indicates the ratio of genes with increased chromatin accessibility upon H_2_O_2_ stress relative to all input genes. The colour scale represents the adjusted p-value of the GO enrichment analysis. The dot size shows the number of genes enriched in this GO term.

Nucleosome-free ATAC-seq peaks in H_2_O_2_-treated and non-treated cells from all three biological replicates were then subjected to principal component analysis (PCA) based on the presence and absence of called peaks. Samples clustered according to their growth conditions, suggesting that the peak calling of the ATAC-seq workflow was able to identify alterations in chromatin accessibility in response to oxidative stress (Figure S3). Notably, the PCA also revealed variations among biological replicates, as replicate three from stressed and non-stressed cells, was separated from the other two replicates by the first principal component (Figure S3, PC1). Hence, we also applied a batch effect correction during differential peak analysis of called ATAC-seq peaks (see Material & Methods for details). In total, 1092 nucleosome-free ATAC-seq peaks with significant signal alterations (FDR < 0.05) in response to oxidative stress were detected (Table S2). Peaks were annotated to the next closest downstream gene to correlate differentially regulated ATAC-seq peaks with transcriptional changes. The majority of genes with increased transcript abundance as suggested by the RNA-seq data set upon H_2_O_2_ treatment also showed elevated chromatin accessibility (Figure 3C). For instance, as already observed by comparing ATAC-seq read coverages between the treatment groups, oxidative stress altered chromatin accessibility of promoter regions and transcript induction of genes including *CAT1, ICL1, OYE32* and *TSA1*, the latter encoding an antioxidative protein ^64^ (Figure 3C). Accordingly, genes downstream of upregulated ATAC-seq peaks represented biological processes involving oxidative stress responses. Interestingly, gene promoters of several genes, including some encoding uncharacterized proteins, displayed significantly elevated chromatin accessibility during H_2_O_2_ treatment without alterations in gene expression. For example, gene C2_09880C, encoding for a putative DNA-binding factor, was among the top 20 genes with increased chromatin accessibility upon oxidative stress. However, no significant alteration in transcription was detected in the corresponding RNA-seq data set (Figure 3C and Table S3). Such cases could represent putative regulators that are primed for transcriptional control in response to additional signaling inputs ^65^. Moreover, it may also reflect the dynamics of a highly transient transcriptional responses often seen in stress adaptation ^54^. In summary, these data demonstrate that ATAC-seq detects changes in chromatin accessibility linked to gene expression control during fungal stress adaptation. Moreover, it can be used to predict novel regulators that are primed for further transcriptional alterations during environmental adaptation.

### Oxidative stress-responsive ATAC-seq peaks are enriched for Cap1 binding sites

Despite assessing the chromatin architecture at different genomic features, ATAC-seq has been used to detect sequence motifs used by transcriptional regulators in ATAC-seq peak regions ^32,39^. This is of special interest for the identification of as yet unknown transcriptional regulators or even regulatory networks governing transcriptional responses. Indeed, regulatory sequences were identified in the malaria parasite *Plasmodium falciparium* during intra-erythrocytic development ^39^. Hence, ATAC-seq can yield new insights into transcriptional networks and their dynamic behavior during environmental adaptations. For example, major virulence traits of *C. albicans*, such as morphological transitions, are under the control of multi-layer genetic regulatory networks engaging both, transcription factors and chromatin modifiers ^20,66,67^. Hence, we tested whether our ATAC-seq data is suitable for motif discovery in *C. albicans*. As mentioned briefly above, the Cap1 transcription factor is the key driver of transcriptional induction during the oxidative stress response ^61^. Its *cis*-acting sequence motif emerged from an earlier study ^45^ (Figure 4A). When we analyzed the enrichment of published Cap1-binding sites in nucleosome-free ATAC-seq peaks upregulated in H_2_O_2_-treated cells, we observed that ATAC-seq read signals were indeed highly increased around the Cap1 motif (Figure 4B). Importantly, a *de novo* motif search identified Cap1 binding sites appearing in ATAC-seq peaks that showed increased intensity after oxidative stress (Figure 4C, motif 3). Of note, some other motifs contained repetitive deoxyadenosine (poly-dA) stretches (Figure 4C, motif 1). While these might be less relevant for the recruitment of transcription factors, such signals can be explained by the preferred occurrence of poly(dA:dT) tracts in nucleosome-depleted regulatory regions, such as TATA elements upstream of the TSS, and the tendency of A-rich elements in TATA-less promoters in *S. cerevisiae* ^68,69^. In addition, GGTTT/AAACC motifs are overrepresented in ATAC-seq data due to a binding bias the Tn5 transposase to such regions ^70^. These data suggest that a more stringent correction of the Tn5 sequence bias is required for a *de novo* motif discovery with higher precision. Notably, tools for advanced sequence bias correction of enzymatically-prepared sequencing libraries, including DNase-seq, MNase-seq and ATAC-seq, are available ^70,71^.

**Figure 4.**
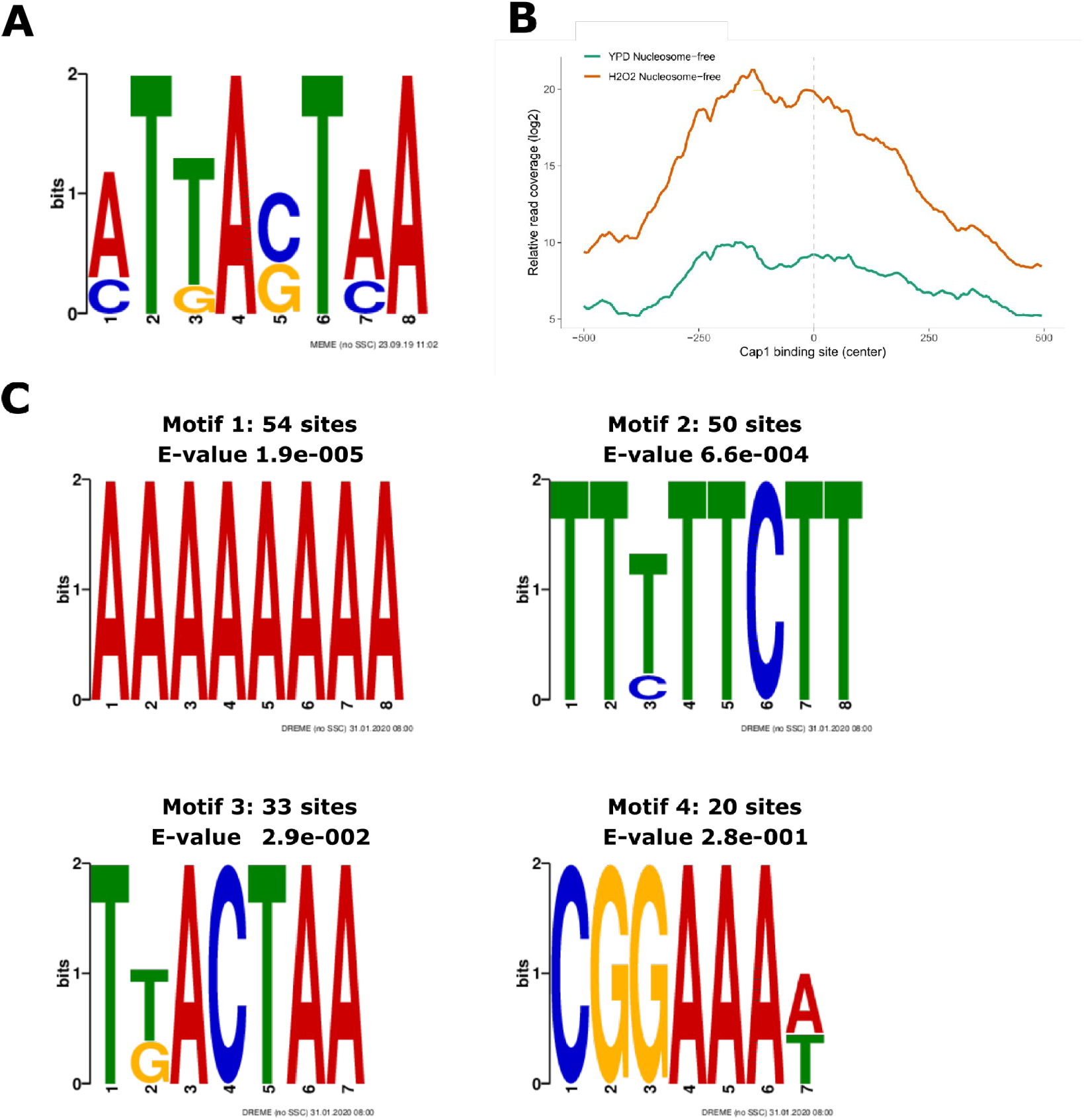
*De novo* motif search in nucleosome-free ATAC-seq peaks with increased signal in response to oxidative stress. **A**) Previously identified Cap1 motif ^45^ used for enrichment analysis in nucleosome-free ATAC-seq peaks with increased signal in H_2_O_2_-treated cells. **B**) Cap1 binding sites are enriched for nucleosome-free ATAC-seq fragments in response to H_2_O_2_. Published Cap1 binding sites ^45^ were used for enrichment analysis within nucleosome-free ATAC-seq peaks. Read coverage in H_2_O_2_-treated and untreated cells around the Cap1 binding sites is shown. **C**) Logos of position weight coverage matrices from *de novo* identified motifs occurring in nucleosome-free ATAC-seq peaks in H_2_O_2_-treated cells (see Material & Methods for details). The number of positive matches among the input sequences and the e-value from the MEME suite DREME tool are indicated above each logo. The top 4 logos are presented.

In conclusion, we here present an integrative workflow for ATAC-seq in the human fungal pathogen *C. albicans*. ATAC-seq offers a new robust tool to capture temporal changes in chromatin landscapes that are tightly connected to transcript abundance during fungal adaptation to stress. In addition, novel regulators, that are not subject to transcriptional control at the time of sample collection, might be identified based on chromatin accessibility signatures. This might be of special interest to capture a broader picture of highly dynamic cellular adaptations where sample drawing is limited. Finally, the combination of ATAC-seq with additional data sets has the potential to predict yet unknown regulatory sequences and transcription factor binding sites, which dictate transcriptional reprogramming and thus, may be crucial for fungal survival upon stress encounter. Finally, the present workflow might be applicable to *in vivo* ATAC-seq approaches, since we were already able to obtain meaningful ATAC-seq data from as little as 50 000 *C. albicans* spheroblasts (data not shown). The re-isolation of comparable fungal cell numbers from host model systems, such as mice, are feasible and thus, ATAC-seq facilitates studies that elucidate fungal responses and adaptation to host defence in a tissue-specific manner. Such data will help to better understand fungal tissue tropism, but also offer to identify new fungal genes amenable to therapeutic intervention.

## Material & Methods

### Fungal strains, media and growth conditions

The *C. albicans* MTL a/α clinical isolate SC5314 ^72^ was used for all experiments and was routinely grown in YPD (1% yeast extract, 2% peptone, 2% glucose; all BD Biosciences) at 30 °C.

To induce oxidative stress, *C. albicans* was first grown overnight to an OD_600_ of approximately one in YPD at 30°C and was then treated with 1.6 mM H_2_O_2_ (Sigma-Aldrich) for 15 minutes while shaking at 30 °C. As control, cells without H_2_O_2_ treatment were cultured in parallel. The cell pellets from these cultures were harvested and directly subjected to spheroblasting and tagmentation (see below).

### Nuclei and genomic DNA (gDNA) isolation, tagmentation, ATAC-seq libraries and sequencing

Nuclei preparation for tagmentation from YPD +/- H_2_O_2_ *C. albicans* was based on a previously published protocol used for *S. cerevisiae* ^31^. Briefly, 3×10^7^ *C. albicans* cells were washed 1x in Sorbitol buffer (1.4 M Sorbitol, 40 mM HEPES-KOH pH 7.5, 0.5 mM MgCl_2_ 10 mM DTT; all Sigma-Aldrich). The pellet was then resuspended in 330 μL Sorbitol buffer and 192 μL of this suspension were mixed with 8 μL of 50 mg/ml 100T Zymolase (2 mg/ml final concentration; Sigma-Aldrich) and incubated for 5 minutes at 30 °C. Spheroblasting efficiency was monitored by diluting a 10 μL aliquot of the spheroblasting reaction into 1 ml distilled water and OD_600_ measurement. After 5 minutes of spheroblasting, the OD_600_ dropped by > 90 % with respect to the initial OD_600_. To sustain oxidative stress for H_2_O_2_-treated cells, 1.6 mM H_2_O_2_ was added to the spheroblasting reaction. Hence, the total time of H_2_O_2_ exposure was 20 minutes. Spheroblasts were then harvested for 5 minutes at 2000 g at 4 °C and washed 1x in ice cold Sorbitol buffer without DTT. The pellet was resuspended in 500 μL ice cold Sorbitol buffer and spheroblasts were counted on a CASY cell counter. Five million spheroblasts were transferred into a tube on ice, harvested 2 minutes at 2000 g at 4 °C and the resulting cell pellet was used for subsequent tagmentation.

#### gDNA isolation

To control for sequence bias by the Tn5 transposase (TDE1) ^46^, gDNA was isolated via phenol-chloroform-isoamyl alcohol (PCI; Sigma-Aldrich) extraction from stationary-phase cells from *C. albicans* as described earlier ^73^ with slight modifications. Briefly, cells were broken up in lysis buffer ^73^ using a FastPrep instrument (MP Biomedicals; settings: 2 rounds of 45 seconds 6 m/s). After PCI extraction, gDNA was precipitated with 100 % ethanol, treated with 10 mg/ml RNase A (Sigm-Aldrich) and again precipitated with ammonium acetate and 100 % ethanol. The final DNA pellet was resuspended in 50 μl TE (10 mM Tris-HCl pH 8, 1 mM EDTA) and subjected to the ATAC-seq library preparation workflow described below.

#### Tagmentation

Five million fungal spheroblasts or 0.5 ng naked gDNA were resuspended in the tagmentation reaction mix (12.50 μL Nextera 2xTD buffer, 2.00 μL Nextera TDE1 [all Illumina], 0.50 μL 50x protease inhibitor cocktail [Roche], 0.25 μL 1 % Digitonin [New England Biolabs] and 10.25 μL nuclease-free distilled H_2_O [ThermoFisher Scientific]) and incubated at 37°C for 30 minutes. The tagmentation reaction was immediately purified using a Qiagen MiniElute PCR purification kit and elution was done in 12 μL elution buffer provided by the kit.

#### ATAC-seq library amplification and size selection

ATAC-seq library preparation was based on the protocol published by the Greenleaf lab ^47^ with minor modifications. Briefly, prior PCR amplification of the ATAC-seq libraries, a test qPCR was performed to determine the optimal number of amplification cycles in order to avoid size and GC bias of the ATAC-seq library ^47^. The qPCR contained the following components: 1 μL tagmented DNA, 0.5 μL Nextera Index primer 1 noMX and Index primer 2.1. barcode (25 μM each), 0.1 μL 100x SYBR green (Sigma-Aldrich, freshly diluted from a 10 000x stock), 5 μL 2x NEBnext High-Fidelity PCR Master Mix (New England Biolabs) and 2.9 μL nuclease-free distilled H_2_O (ThermoFisher Scientific). The qPCR was performed in a Realplex Mastercycler (Eppendorf) with the following cycling conditions: 72 °C 5 minutes, 98 °C 30 seconds, 25 cycles of 98 °C 10 seconds, 63 °C 30 seconds, 72 °C 1 minute and a final hold step at 10 °C. The cycle number for optimal ATAC-seq library amplification was determined as in ^47^. The enrichment PCR was then performed using 10 μL tagmented DNA, 2.5 μL Index primer 1 noMX and Index primer 2.x. barcode (25 μM each), 25 μL 2x NEBnext High-Fidelity PCR Master Mix (New England Biolabs) and 10 μL nuclease-free distilled H_2_O (ThermoFisher Scientific). The PCR reaction was then incubated in a PCR thermocycler using the following conditions: 72 °C 5 minutes, 98 °C 30 seconds, x cycles (depending on the test qPCR results) of 98 °C 10 seconds, 63 °C 30 seconds, 72 °C 1 minute and a final hold step at 10 °C. All ATAC-seq libraries from this study were PCR amplified using 11 cycles. For multiplexing, a different Nextera Index primer 2 barcode was used for each sample. See Table S1 for a list of Nextera PCR primers used for library amplification in this study.

The amplified ATAC-seq libraries were immediately purified and size selected with a double-sided solid-phase reversible immobilization (SPRI) approach (0.5x/1.4x) using AMPure XP beads (Beckman Coulter) according to the manufacturer’s instructions. Final DNA elution was done with elution buffer provided in the Qiagen MiniElute PCR purification kit. The final ATAC-seq libraries were quantified using the fluorescent dye Quantifluor (Promega, Vienna, Austria) according to the manufacturer’s manual. The yield for each ATAC-seq library sample was between 60 and 90 nM for replicate 1-2 and around 13 nM for replicate 3.

The quality of purified libraries was analyzed on a Bioanalyzer High Sensitivity DNA chip (Agilent; see Figure 1 for an example) by following the manufacturer’s instructions.

#### Next generation sequencing

ATAC-seq libraries were prepared from three biological replicates for each condition (YPD, YPD+H_2_O_2_ and gDNA) and pooled in equimolar ratios for sequencing. Sequencing was done in 75bp paired-end read mode on a HiSeq 4000 system at the Biomedical Sequencing Facility (BSF; https://cemm.at/research/facilities/biomedical-sequencing-facility-bsf/) at the Center of Molecular Medicine (CeMM) in Vienna, Austria.

### ATAC-seq data analysis workflow

#### Pre-processing and read alignment

Quality control of raw sequencing reads (bam) was done using fastQC v0.11.8 ^74^. Illumina TrueSeq adapter trimming was done via cutadapt v1.18 (https://cutadapt.readthedocs.io/en/stable/; settings: --interleaved -q 30 -O 1). Trimmed reads were then aligned to the haploid *C. albicans* SC5314 genome (assembly 22, version A22-s07-m01-r88; http://www.candidagenome.org/) using NextGenMap v0.5.5 ^75^ with only keeping aligned reads with a minimum mapping quality of 30 (settings: -b -p -Q 30). Optical read duplicates were removed using Picard tools (Broad Institute, https://broadinstitute.github.io/picard/, settings: MarkDuplicates REMOVE_DUPLICATES=true VALIDATION_STRINGENCY=LENIENT; Broad Institute, https://broadinstitute.github.io/picard/) and mitochondrial reads were removed using the ‘intersect’ tool from BEDTools with −v settings (https://github.com/arq5x/bedtools2). The average number of mapped reads per conditions for each chromosome and the size of each chromosome as comparison are shown in Figure S1. Chromosome sizes of *C. albicans* were retrieved from the National Center for Biotechnology Information (NCBI). The ATAC-seq fragment length distribution from properly paired reads is represented in Figure 1C.

Aligned *bam* files were split according to the fragment lengths of the sequencing read pairs as done previously ^30^. Read fragments below 100 bp were considered coming from nucleosome-free regions and were used for further analysis. Normalized read coverage files (bigwig) of nucleosome-free read fragments were created using deepTools2 ‘bamCoverage’ (^76^; settings: -e -bs 5 --normalizeUsing CPM) and visualized using the Integrative Genomics Viewer (IGV; ^77^).

#### Prediction of nucleosmal positions and genomic coverage analysis of ATAC-seq signals

Nucleosomal occupancies were predicted using NucleoATAC ^31^ with default parameters. NucleoATAC analysis was performed for all *C. albicans* promoter regions (TSS -/+ 1000 bp). Promoter regions were extracted using the ‘promoters’ function from the Bioconductor GenomicRanges package ^78^. The NucleoATAC bedgraph output files were further converted to the bigwig format using the bedGraphToBigWig tool (https://github.com/ENCODE-DCC/kentUtils). The NucleoATAC result was further compared with the nucleosome-free ATAC-seq reads (see above) and published MNase-seq data for *C. albicans* grown in YPD ^49^. MNase-seq raw data sets were downloaded via sequence read archive (SRA) and processed as the ATAC-seq raw data. The aligned bam files were converted into a normalized read coverage (bigwig) file using the deepTools2 ‘bamCoverage’ function (^76^, settings: as above, except −bs 1). Genomic read coverage tracks were visualized with the IGV. To plot the read coverage of nucleosome-free ATAC-seq peaks, nucleosome positions from the NucleoATAC analysis and the published MNase-seq data ^49^ over all *C. albicans* transcripts, a coverage matrix was computed using the deeptools2 ‘computeMatrix’ function (^76^; reference-point -a 1000 -b 1000 -bs 5). Transcripts from *C. albicans* were extracted by the ‘transcripts’ function of the GenomicRanges package ^78^ using the current genomic annotation from the *C. albicans* assembly 22 (version A22-s07-m01-r88; http://www.candidagenome.org/). MNase-seq reads were extended to 100 bp prior coverage plotting as described in the original publication ^49^.

Differential read coverage (log2-ratio) in nucleosome-free ATAC-seq peaks between H_2_O_2_-treated and non-treated samples was analyzed by the deeptools2 ‘bamCompare’ function (^76^; settings: --normalizeUsing CPM -bs 5 -e --scaleFactorsMethod None). For downstream analysis, this output was first used to compute a coverage matrix with the deeptools2 ‘computeMatrix’ tool ^76^ as described above across promoter regions (−1000/+200bp with respect to the TSS) of all *C. albicans* transcripts. Heatmap and coverage plots were generated using the ‘plotHeatmap’ function from deeptools2 ^76^ using k-means clustering (settings: -kmeans 4). Coverage regions from cluster 1 and cluster 4 were further subjected to GO term analysis using the ‘enrichGO’ function from the clusterProfiler package (^79^; settings: ont = “BP”, qvalueCutoff = 0.05, readable = TRUE). The GO term analysis result was plotted using the clusterProfiler ‘dotplot’ and ‘cnet’ function ^79^.

#### Peak calling and genomic annotation of ATAC-seq peaks

Peak-calling for each individual sample was done with MACS2 v2.1.2 using ‘callpeak’ (^80^; settings: -f BAMPE -g 14521502). The aligned read fragments from the gDNA ATAC-seq libraries were merged into one bam file using the samtools ‘merge’ function ^81^ and used as a background control for peak calling. Peaks from all samples and replicates were merged and converted to the gff format for read counting using htseq-count (^82^; settings: -f bam -s no -t peak) and differential ATAC-seq peak analysis (see below). The reproducibility of called ATAC-seq peaks among biological replicates was analyzed via principal component analysis (PCA) using the ‘prcomp’ function from the R stats package (https://www.rdocumentation.org/packages/stats) and is presented in Figure S3. For the ATAC-seq peak annotation, the genomic annotation from *C. albicans* assembly 22 (version A22-s07-m01-r88) was downloaded from the CGD (http://www.candidagenome.org/) and was merged with the 5’UTR and TSS annotations of *C. albicans*, which were retrieved from the yeast transcription start site (YeasTSS) database (http://www.yeastss.org/). This file was further used to create a TxDb object using the GenomicFeatures ‘makeTxDbFromGFF’ function (^78^; settings: dataSource = “CGD”, organism = “Candida albicans SC5314”, circ_seqs = “Ca22chrM_C_albicans_SC5314”). Peak annotation was done using the ChIPseeker Bioconductor package ^83^ with the ‘annotatePeak’ function (options: tssRegion = c(−2000, 0), genomicAnnotationPriority = c(“Promoter”, “5UTR”, “3UTR”, “Exon”, “Intron”, “Downstream”, “Intergenic”)). The distribution and overlaps of peak annotations among genomic features was visualized with the ‘upsetplot’ function from the ChIPseeker package. The ATAC-seq peak location across all chormosomes and the fold change in peak signals between H_2_O_2_ treated and non-treated samples were visualized using the karyoploteR package ^84^.

#### Analysis of differential ATAC-seq peaks

Differential accessible peak detection was done using the edgeR package ^85^ with the generalized linear model (glm) approach. Since replicate three clustered away from the other two replicates in the PCA analysis (Figure S3), the glm approach was combined with batch effect correction. Table S2 contains a complete list of detected ATAC-seq peaks and the edgeR analysis result. GO term analysis of genes with significantly increased peak signal (FDR < 0.05) in upstream regions upon H_2_O_2_ stress was performed with the ‘enrichGO’ function from the clusterProfiler package ^79^ as described above.

#### Motif search

Nucleosome-free ATAC-seq peaks with increased signal in the H_2_O_2_-treated samples (FDR < 0.05) from the edgeR analysis were further subjected to motif search analysis using the MEME suite (v5.1.0; ^86^; http://meme-suite.org/). First, the previously identified *C.albicans* Cap1 regulon ^45^ was used to generate the position weight matrix for Cap1 with the MEME web application (settings: 1 occurrence per sequence, motif width = 8). This was further used as input for the MEME FIMO tool ^87^ to analyze Cap1 binding sites in the upregulated nucleosome-free ATAC-seq peaks in H_2_O_2_-treated samples (settings: --parse-genomic-coord). The enrichment of the identified Cap1 binding sites was re-formatted to bed format and further used to compute a coverage matrix with the ‘computeMatrix’ function from deeptools2 (^76^; settings: reference-point, -b 500 -a 500 -bs 5 --referencePoint center) in nucleosome-free ATAC-seq peaks from H_2_O_2_-treated and non-treated samples. For *de novo* motif analysis, upregulated nucleosome-free ATAC-seq peaks during oxidative stress were further filtered based on their occurrence within the 1 kb upstream region of a gene and a maximum peak width of 500 bp. The resulting 184 regulated peaks were used for *de novo* motif search using the online version of the MEME suite tool DREME (^88^; parameters: E-value threshold 1; search given strand only). Figure 4C shows the top four identified motifs.

#### Data plotting

Unless otherwise stated, plots were done with the ‘ggplot2’ package in R ^89^.

### External data sets

Our previously published (^116^) RNA-seq data from *C. albicans* in response to 15 minutes H_2_O_2_ treatment (Gene Expression Omnibus [GEO] accession number GSE73409) were used to compare ATAC-seq peaks with increased signal in H_2_O_2_-treated samples with genes upregulated in response to oxidative stress. MNase-seq data (^449^; described above) was downloaded from SRA (SRR059732).

## Code availability

The entire bioinformatics analysis pipeline is freely accessible on github: https://github.com/tschemic/ATACseq_analysis

## Data availability

ATAC-seq data will be deposited at the GEO server upon manuscript publication.

## Acknowledgements

This work was funded by the grants *ChromFunVir* (Project FWF-P-31712-B21) and FWF-SFB-070-HIT from the Austrian Science Fund to KK. MT was supported by an Erwin Schroedinger Fellowship (J3835) of the Austrian Science Fund. We are grateful to the Christoph Bock lab members Thomas Krausgruber and André Rendeiro for advice about the ATAC-seq sample preparation and bioinformatics analysis. In addition, we want to acknowledge Aaron Hernday and Mohammad Qasim for helpful and constructive discussion about the initial set up of ATAC-seq in *C. albicans*.

## Author contributions

S.J., M.T. and K.K. designed the research. S.J. and T.M. performed experiments. M.T. and S.J. analyzed data. M.T. established the bioinformatics workflow. S.J. and K.K wrote, revised and edited the manuscript. M.T. revised and edited the manuscript.

