## Supplemental Information for "ATAC-seq identifies chromatin landscapes linked to the regulation of oxidative stress in the human fungal pathogen *Candida albicans*"

From the
[1]Department of Medical Biochemistry, Max Perutz Labs Vienna, Medical University of Vienna, Campus Vienna Biocenter, Dr. Bohr-Gasse 9/2, A-1030 Vienna

Running title: ATACing fungal chromatin

* contributed equally
### To whom correspondence should be addressed


**Figures:**

**Figure S1:** Number of mapped ATAC-seq reads to each *C. albicans* chromosome.

**Figure S2:** Analysis of genes with decreased chromatin accessibility in regions upstream their TSS.

**Figure S3:** PCA of detected nucleosome-free ATAC-seq peaks.

**Tables:**

**Table S1:** Nextera primers used for ATAC-seq library preparation

**Table S2:** MACS2 peak calling and edgeR analysis result.

**Table S3:** Genes with increased chromatin accessibility upon oxidative stress without alterations in gene expression.

#### Figure S1

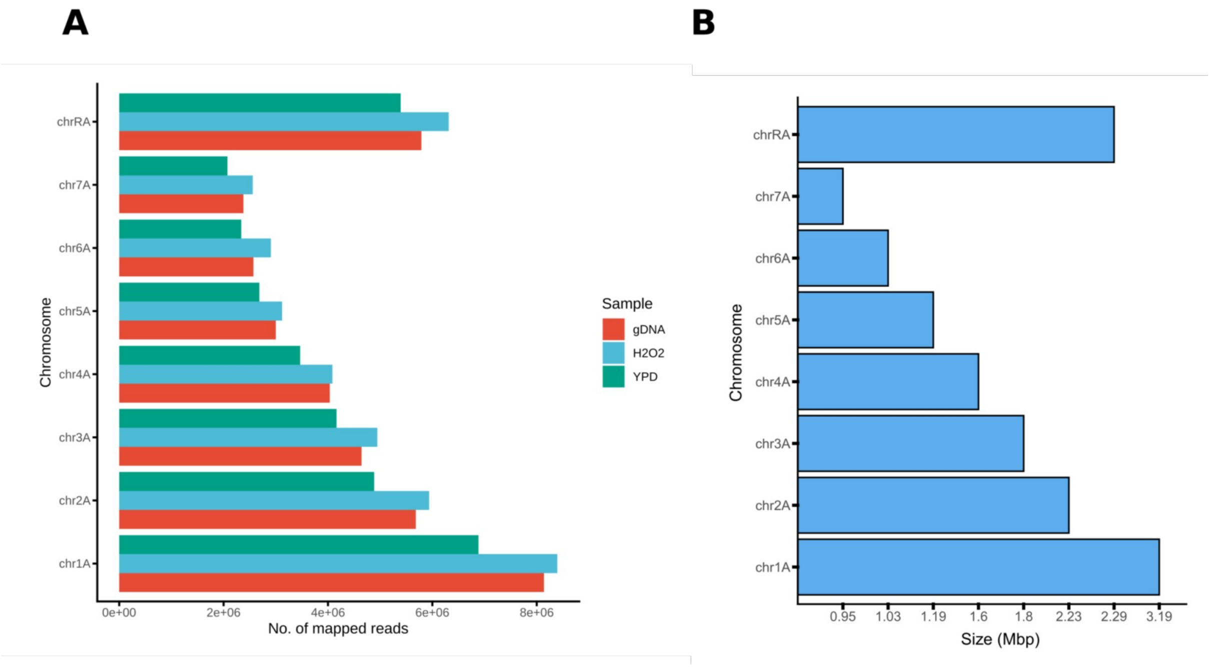


**Figure S1. Number of mapped ATAC-seq reads to each *C. albicans* chromosome.**
**A-B)** The aligned reads from each biological replicate of the gDNA, $H_2O_2$-treated (H2O2) and non-treated (YPD) ATAC-seq libraries were merged and the number of mapped reads (x-axis) per chromosome (y-axis) was plotted (A). As comparison, the total size (in Mbp) of each *C. albicans* chromosome is presented as well (B).

### Figure S2

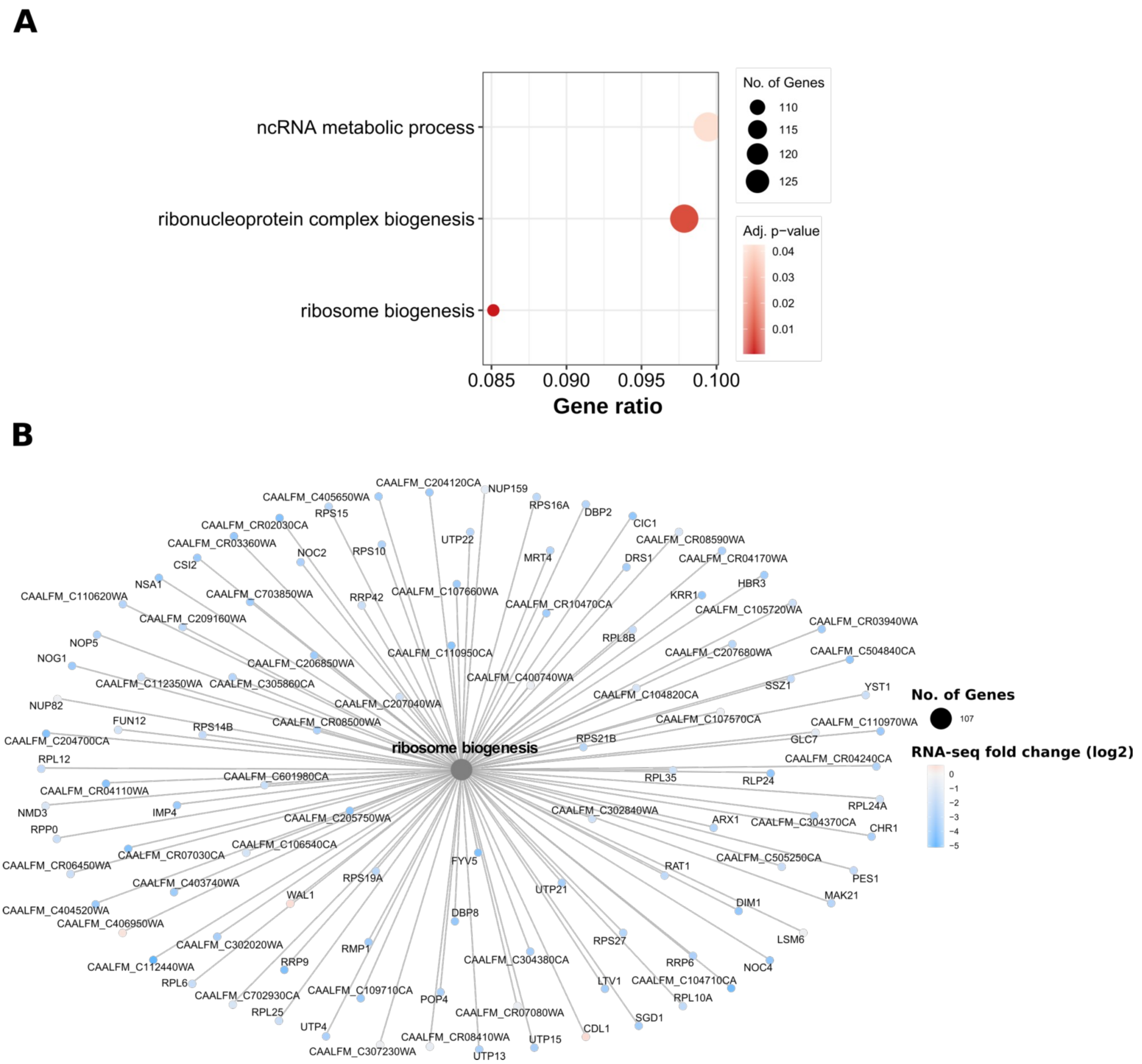


**Figure S2. Analysis of genes with decreased chromatin accessibility in regions upstream their TSS.** **A-B)** Genomic regions from cluster 4 (see Figure 2C-D) were annotated to the next downstream gene and subjected to GO term enrichment analysis. Enriched biological processes are represented as dot plot (A) and cnet plot for the biological process “ribosome biogenesis” (B). See figure legend from Figure 2C-D for detailed description.

### Figure S3

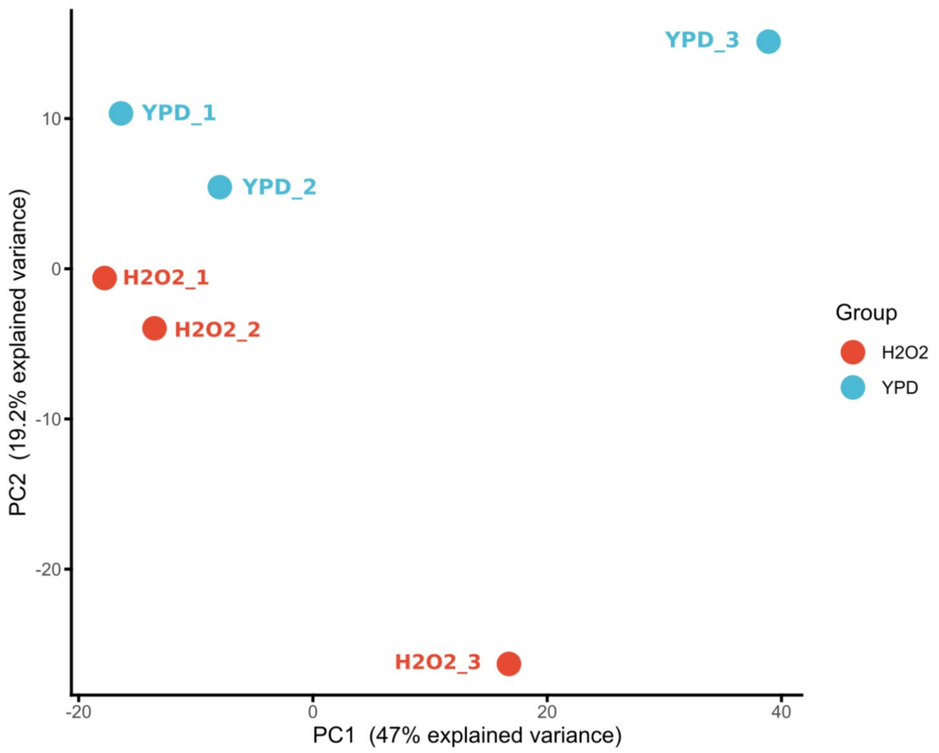


**Figure S3. PCA of detected nucleosome-free ATAC-seq peaks.**
Called nucleosome-free ATAC-seq peaks from all samples were merged and subjected to PCA based on the presence or absence of ATAC-seq peaks in each replicate and condition (YPD and $H_2O_2$). Each dot in blue represents three biological replicates from YPD-grown cells (YPD) and each dot in red represents three biological replicates of $H_2O_2$-treated samples (H2O2). The x-axis shows the principal component 1 (PC1) and the y-axis the principal component 2 (PC2).

**Table S1. Nextera primers used for ATAC-seq library preparation.**

| name | barcode | sequence (5' -> 3') |
|---|---|---|
| Ad1_noMX | | AATGATACGGCGACCACCGAGATCTACACTCGTCGGCAGCGTCAGATGTG |
| Ad2.30_TTGCTAAG | TTGCTAAG | CAAGCAGAAGACGGCATACGAGATCTTAGCAAGTCTCGTGGGCTCGGAGATGT |
| Ad2.31_ATAAGTTA | ATAAGTTA | CAAGCAGAAGACGGCATACGAGATTAACTTATGTCTCGTGGGCTCGGAGATGT |
| Ad2.32_ATCACTCG | ATCACTCG | CAAGCAGAAGACGGCATACGAGATCGAGTGATGTCTCGTGGGCTCGGAGATGT |
| Ad2.33_GTTAACAG | GTTAACAG | CAAGCAGAAGACGGCATACGAGATCTGTTAACGTCTCGTGGGCTCGGAGATGT |
| Ad2.38_GAAATGCC | GAAATGCC | CAAGCAGAAGACGGCATACGAGATGGCATTTCGTCTCGTGGGCTCGGAGATGT |
| Ad2.39_AACGCCAT | AACGCCAT | CAAGCAGAAGACGGCATACGAGATATGGCGTTGTCTCGTGGGCTCGGAGATGT |
